# Prediction of GPI-Anchored proteins with pointer neural networks

**DOI:** 10.1101/838680

**Authors:** Magnús Halldór Gíslason, Henrik Nielsen, José Juan Almagro Armenteros, Alexander Rosenberg Johansen

**Author notes:** Corresponding author (H. Nielsen).

## Abstract

GPI-anchors constitute a very important post-translational modification, linking many proteins to the outer face of the plasma membrane in eukaryotic cells. Since experimental validation of GPI-anchoring signals is slow and costly, computational approaches for predicting them from amino acid sequences are needed. However, the most recent GPI predictor is more than a decade old and considerable progress has been made in machine learning since then. We present a new dataset and a novel method, NetGPI, for GPI signal prediction. NetGPI is based on recurrent neural networks, incorporating an attention mechanism that simultaneously detects GPI-anchoring signals and points out the location of their *ω*-sites. The performance of NetGPI is superior to existing methods with regards to discrimination between GPI-anchored proteins and other secretory proteins and approximate (±1 position) placement of the *ω*-site.

NetGPI is available at: https://services.healthtech.dtu.dk/service.php?NetGPI

The code repository is available at: https://github.com/mhgislason/netgpi-1.1

## 1. Introduction

Some of the proteins that follow the secretory pathway are bound to the membrane of eukaryotic cells by specific mechanisms. One of these mechanisms is a post-translational modification where a glycosylphosphatidylinositol (GPI) anchor is attached to the protein. The identification of proteins that undergo this modification is of high interest due to the diversity of functions that they perform. GPI-anchored proteins are essential in the development of fungi and animal cells [3, 13]. They are also involved in certain diseases such as paroxysmal nocturnal haemoglobinuria, an acquired haematopoietic stem-cell disorder [31], and in the defense mechanisms of various protozoan parasites such as *Leishmania* and *Trypanosoma* [17]. Consequently, the development of computational tools that are able to detect proteins with this modification is of high impact on the research of eukaryotic cell biology [18].

GPI-anchored proteins have two signals in their primary sequence: an N-terminal sequence for endoplasmic reticulum targeting (signal peptide) and a C-terminal signal sequence directing the attachment of the GPI-anchor. This attachment is carried out by a GPI transamidase which recognizes the C-terminal signal sequence and cleaves the peptide bond at the GPI-anchor attachment site, known as the *ω*-site. This cleavage creates a covalent bond between the GPI and the C-terminus of the cleaved protein, allowing the protein to remain tethered to the membrane. C-terminal signal sequences are generally composed of five regions, which are determined by the amino acids before the *ω* site (*ω*-minus) and after (*ω*-plus). The five regions are: a stretch of polar amino acids that form a flexible linker region (*ω* − 10 to *ω* − 1); the *ω* site amino acid; the *ω* + 2 amino acid, a restrictive position with mostly G, A, or S; a spacer region of moderately charged amino acids (*ω* + 3 to *ω* + 9 or more), and a stretch of hydrophobic amino acids starting approximately at *ω* + 10 [23].

In order to detect proteins that carry this signal, experimental assays are required. Such experiments are generally low throughput and costly, which has resulted in a low amount of experimentally annotated GPI-anchored proteins. To overcome this limitation, fast computational methods that can approximate the experimentally validated process are needed. For this purpose, current machine learning methods exist for predicting GPI-anchors [6, 9, 26]. However, these methods were developed more than a decade ago and do not utilize recent progress in machine learning methods nor access to new data sources. Deep learning methods, such as recurrent neural networks (RNN) [10], have recently proven effective at protein prediction tasks [12]. However, deep learning requires large amounts of annotated samples to generalize well [15].

In this paper we present a new tool for detecting GPI-anchored proteins and determining the position of the *ω*-site using recurrent neural networks. To overcome the low amounts of experimentally validated data we build a new dataset composed of both experimentally annotated and predicted GPI anchored proteins. To benchmark our method against previous methods, we only consider experimentally annotated samples. Regardless, our method achieves state-of-the-art performance on the GPI-anchor prediction task. Moreover, we show that the model learns biologically meaningful characteristics.

### 1.1. Related works

Initial work on predicting the presence of GPI-anchors and the *ω*-site was published by Eisenhaber et al. [6]. This work, known as the Big-∏ Predictor, details a method that evaluates amino acid type preferences at positions near a potential *ω*-site as well as the concordance with general physical properties encoded in multi-residue correlation within the motif sequence [6]. Big-∏ provides kingdom-specific predictions as it was trained on metazoan, protozoan, fungi [7], and plant [8] proteins separately.

Fankhauser and Mäser [9] presented a neural network based prediction tool called KohGPI/GPI-SOM. GPI-SOM utilizes a Kohonen Self Organizing Map structure, which takes as input the average position of a given amino acid relative to its proximity to the C-terminal, the hydrophobicity of the amino acid at 22 C-terminal positions, and 2 units representing the quality of the presumed *ω*-site and its position. Both GPI-SOM and Big-∏ utilize an external signal peptide predictor known as SignalP [1] to preselect proteins.

FragAnchor was published by Poisson et al. [27] in 2007. FragAnchor uses a feed-forward neural network model to detect potential GPI-anchoring signal sequences and a Hidden Markov Model (HMM) to quantify the prediction confidence and to estimate the position of the *ω*-site, the spacer region and the hydrophobic tail. Like the previous two methods, FragAnchor relies on external evidence for the signal peptide and only regards the last 50 C-terminal amino acids. Unfortunately the prediction tool is no longer available online.

In 2008 Pierleoni, Martelli & Casadio published Pred-GPI, a GPI-anchor predictor using a Support Vector Machine (SVM) for the GPI-anchoring signal discrimination and a HMM to predict the position of the *ω*-site [26]. The HMM has 46 states with varying probabilities for amino acids and the potential *ω*-site assigned to the 26th state. The SVM takes as input the negative log-likelihood computed by the HMM as well as 82 features intended to describe the overall composition of the sequence, the features of the N-terminal regions comprising the signal peptide, and the features of the C-terminal regions containing the cleaved GPI-anchor signal. Pred-GPI supplies two different variants: one model where the potential *ω*-site is restricted to be one of Cysteine, Aspartic acid, Glycine, Asparagine, and Serine – this approach they refer to as the conservative model – and a non-conservative variant which has no such restriction. Unlike the other three methods, Pred-GPI does not rely on an external signal peptide predictor, such as SignalP.

## 2. Materials and methods

### 2.1. Dataset

All data used in this project are extracted from the UniProt database, release 2019_02 [32]. The dataset construction follows two main steps: data gathering and homology partitioning. First, we select 3618 eukaryotic proteins found with experimental evidence (ECO:0000269) of a signal peptide. All proteins are truncated to the last 100 amino acid positions. This is because our method does not include the prediction of the signal peptide and it is assumed that all relevant sequence information resides in or near the C-terminal positions. Instead, it relies on experimental evidence for the signal peptide or signal peptide prediction tools, such as SignalP [1]. After truncation we remove exact duplicates, leaving 3567 unique sequences. Of the 3567, there are 981 assumed to be GPI-anchored. Out of the 981 there are 161 with experimental evidence for the GPI-anchoring signal of which 50 also have experimental evidence for the *ω*-site, where the GPI-anchor would be attached. The remaining 820 have non-experimental evidence for the presence of a GPI-anchoring signal. This leaves 2586 without any evidence for the presence of a GPI-anchoring signal, which are assumed to be not GPI-anchored. The samples are also labelled by kingdom taxonomy: animal, fungi, plant, or other. The dataset composition is fully detailed in table 1.

**Table 1.**
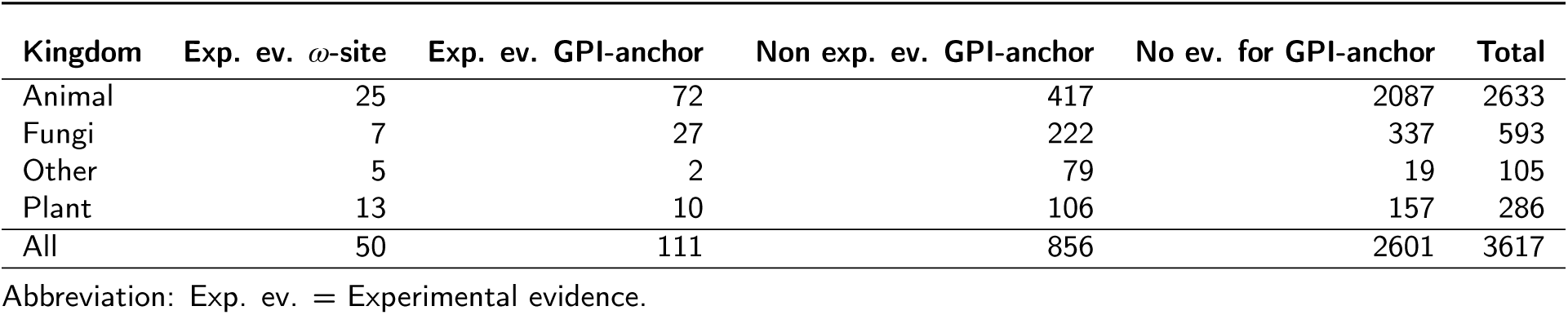
The dataset composition. There are 161 samples in total with experimental evicence for a GPI-anchoring signal. The samples with experimental evidence for the *ω*-site also have experimental evidence for a GPI-anchoring signal, here however, they are presented separately

| Kingdom | Exp. ev. ω-site | Exp. ev. GPI-anchor | Non exp. ev. GPI-anchor | No ev. for GPI-anchor | Total |
| --- | --- | --- | --- | --- | --- |
| Animal | 25 | 72 | 417 | 2087 | 2633 |
| Fungi | 7 | 27 | 222 | 337 | 593 |
| Other | 5 | 2 | 79 | 19 | 105 |
| Plant | 13 | 10 | 106 | 157 | 286 |
| All | 50 | 111 | 856 | 2601 | 3617 |

Abbreviation: Exp. ev. = Experimental evidence.

Homology partitioning is the separation of a set of nucleic or protein sequences into subsets, such that all sequences within each subset are non-homologous to sequences in other subsets. Commonly used clustering tools, such as CD-HIT [16], MMSeqs2 [30] and BLASTCLUST [5], provide fast homology separation, where all sequences within each subset are homologous to one another. However, to achieve fast separation, they employ approximate alignment and separation procedures. These approximations are acceptable when the only goal is to ensure that all samples within each subset are homologous to one another, however, this is at the cost of, potentially, high similarity between samples in different subsets.

To homology partition the dataset we define percent identity of 30% as the threshold. We follow a four phase procedure. First we obtain global alignments, using the program `ggsearch36`, which is a part of the `FASTA` package [25]. The program implements the Needleman-Wunsch algorithm for global alignments [22]. We set the program’s -E parameter, which is the expectation value threshold, to be larger than the dataset size. The percent identity is provided, however the denominator depends on the chosen output format. The default output format calculates percent identity with the length of the alignment as the denominator. In the second phase, we cluster the pairwise percent identities, using restricted single-linkage clustering. As the end-goal is to partition the dataset into five comparable subsets, the clustering procedure is restricted such that no single cluster is allowed to have more than 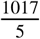 samples labelled GPI-anchored or 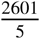 samples labelled not GPI-anchored. In the third phase, the clusters are grouped together into five partitions, such that the number of GPI-anchored and not GPI-anchored samples is comparable across all partitions. In the final phase, samples are removed until no percent identity above 30% is found between samples in different partitions. During the removal phase, samples can be moved between partitions, as an attempt to reduce the number of samples removed. The composition of the final partitions can be seen in table 2. The final dataset contains 966 proteins labelled GPI-anchored and 2573 labelled not GPI-anchored, for a total of 3539. Out of the 28 removed, one has experimental evidence for a GPI-anchoring signal sequence. All 50 samples with an experimentally verified *ω*-site are retained.

**Table 2.** The dataset, partitioned using Needleman-Wunch, global alignment, pairwise percent identity (PID), to 30% PID. The global alignments are obtained using the `ggsearch36` program, provided with the `FASTA` package.

| Partition: | 0 | 1 | 2 | 3 | 4 | Label | Mean | Samples |
| --- | --- | --- | --- | --- | --- | --- | --- | --- |
|  | 188 | 189 | 225 | 169 | 195 | Anchored | 193.2 | 966 |
|  | 488 | 484 | 484 | 629 | 488 | Not anchored | 514.6 | 2573 |

### 2.2. Objective

The objective of GPI prediction is to decide whether a GPI signal is present and, if present, to determine the position of the *ω*-site in a protein sequence. We combine these two tasks by reducing them to the single task of maximizing the probability of a position in a sequence. To achieve this, we add a placeholder to the end of the protein sequence which serves as an indicator for the absence of a GPI-anchoring signal. Thus, we formally define the objective as maximizing the probability of a position in 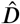, which is known as pointing [33].

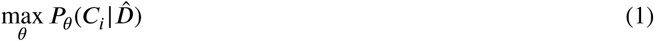

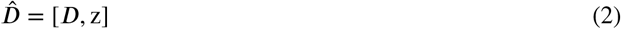

Where *D* ∈ Σ^*T*−1^ is an amino acid sequence and Σ. is a dictionary of the twenty common amino acids as well as the token X, which represents any encountered amino acid not in Σ. We only consider the last 100 amino acids in the protein sequence, such that the length *T* − 1 ≤ 100. If the sequence does not contain an *ω*-site we maximize the probability of the protein being non GPI-anchored. Inspired by work in natural language processing [20, 19], we represent the lack of an *ω*-site by maximizing the placeholder position known as the sentinel, *z*, at the end of the amino acid sequence. This results in 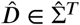 where 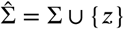. *C*_*i*_ then corresponds to a position in 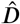.

To parameterize the conditional probability distribution *P*_*θ*_ we use a neural network architecture known as the Long-Short Term Memory (LSTM) Cell [11] and distributed representations of the amino acids [21] as shown in equation 3,

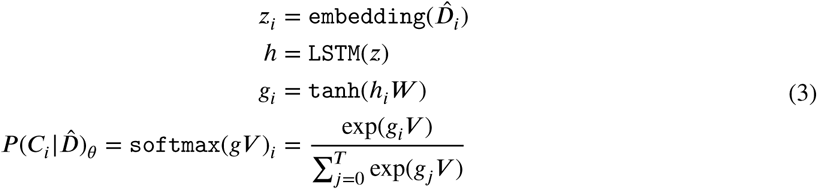

where `embedding` : 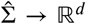 turns each amino acid into a distributed representation of real numbers using a linear trainable weight of size *d* and 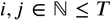 are indexes of the protein sequence including the sentinel position. The `LSTM` is a non-linear transformation of a sequence of real values. It uses trainable recurrent units to distribute sequential information across the protein sequence, `LSTM` : ℝ^*T×d*^ → ℝ^*T×d′*^, where *d*′ is the output size of the `LSTM`. As we use a bidirectional `LSTM` [29] we end up with two hidden representations of size *d*′. To get the probability over the sequence we project the output of every position to a logit, *g*_*i*_*V* ∈ ℝ, followed by a `softmax` : ℝ^*T*^ → [0,1]^*T*^ that normalizes the logits into a probability distribution over the sequence. To create the logits we use a two layer feed forward neural network on top of the `LSTM` hidden states, *h* ∈ ℝ^*T*×2*d′*^, with a `tanh` activation function, *W* ∈ ℝ^*2d′×d′′*^, and *V* ∈ ℝ^*d′′*^. This usage of `softmax` over a sequence is a modification of attention where the interaction size *d*′′ of *gV* is the attention hidden representation size. This modification of attention is known as a pointer network [33].

The `embedding, LSTM`, *W*, and *V* are all trainable with stochastic gradient descent using back-propagation through time [34]. We have visualized our model in Figure 1.

**Figure 1:**
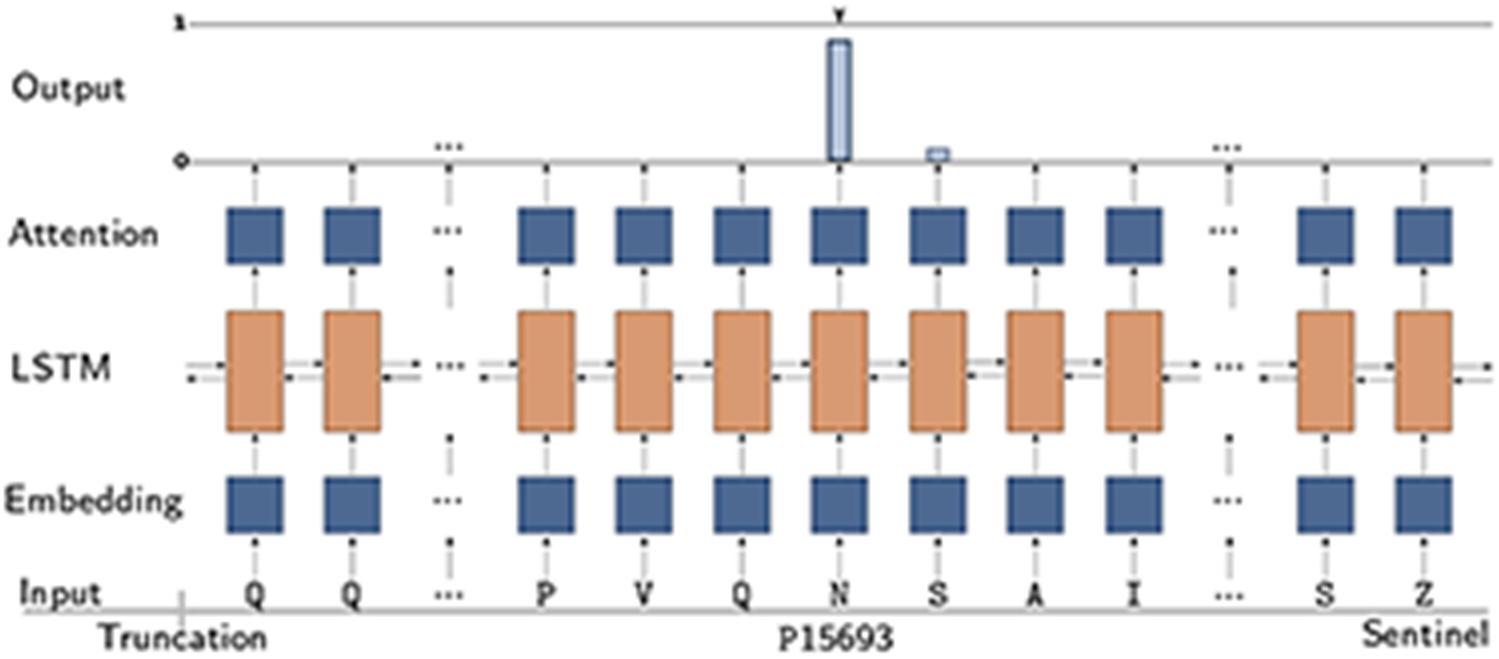
Diagram of the model, illustrating how the model points to a position in a sequence, in this case, the entry with UniProt accession number P15693. The sequence is truncated to the last 100 amino acids and the sentinel, *z*, is appended. The predicted *ω*-site is an Asparagine (N). If the position with highest likelihood had been the sentinel position, then the protein would have been predicted as non GPI-anchored.

### 2.3. Model Details

All partitions contain samples, which will be used for testing, therefore five-fold nested cross-validation is used for a generalized estimate of the performance of potential hyperparameter combinations and to test the performance of the final selection. In all, there are 20 models trained. For each partition, the four other partitions are cross-validated where three partitions are used to train a model and one to validate the performance. To compare the performance of different combinations of hyperparameters, within a validation group, the average performance of the four models, is compared with the average performance using other hyperparameter combinations. The `SIGOPT` platform is used for model selection [4]. The `SIGOPT` optimization engine provides suggestions for hyperparameters based on inference from the performance of other hyperparameter combinations, measured with any real valued metric. The user supplies the values and specifies if the value should be minimized or maximized. Each of the five sets of four models is trained with at least 200 hyperparameter combinations. The hyperparameters to tune are: The size of the distributed representation (*d*), the LSTM cell hidden representation (*d*′), the number of LSTM layers, the LSTM dropout, the attention hidden representation (*d*′′), the batch size, and the optimizer’s learning rate, learning rate decay and weight decay. Each epoch, after the first, the learning rate is updated by multiplying it with the learning rate decay. A model is then trained for each cross-validation split, on a shortlist of hyperparameter combinations, shown in table 3. Each combination is executed 30 times and the model with the highest validation performance is used as the final model. Each set of four models is used as an ensemble predictor for the respective test partition. The logarithm of the probability distribution of each of the four models is averaged and used as an ensemble prediction. The web service predictions are generated with a 20-model ensemble. The neural network is trained with stochastic gradient descent using the Adam optimizer [14]. The models are implemented with the PyTorch deep learning framework [24].

**Table 3.** Four combinations of hyperparameters with the best validation performance, every model has 4 `LSTM` layers and is trained for 300 epochs.

| e.d. (*d*) | LSTM h.u. (*d'*) | LSTM dropout | a.d. (*d''*) | lr. | lr. decay | w. decay | batch size |
| --- | --- | --- | --- | --- | --- | --- | --- |
| 16 | 20 | 0.62 | 340 | 0.002479 | 0.9970 | 0.001930 | 32 |
| 16 | 16 | 0.55 | 283 | 0.003346 | 0.9987 | 0.009095 | 128 |
| 16 | 16 | 0.6 | 283 | 0.003346 | 0.9987 | 0.010052 | 128 |
| 22 | 22 | 0.6 | 283 | 0.003346 | 0.9987 | 0.009095 | 128 |

Abbreviations: e.d. = embedding dimension, h.u. = hidden units, a.d. = attention dimension, lr. = learning-rate, w. = weight.

#### 2.3.1. Quantitative evaluation criteria

To evaluate the discrimination between GPI-anchored and non GPI-anchored proteins we use the Matthews Correlation Coefficient (MCC) and for *ω*-site prediction evaluation we use the F1 score [2]. The F1 score is the harmonic mean of sensitivity (how many of the true cleavage sites are predicted correctly) and precision (how many of the predicted cleavage sites are true). Due to the dual nature of the problem, and the lack of experimental *ω*−site evidence in the training set, a simple heuristic is devised. The heuristic is a composition of the two evaluation methods. The F1 score is calculated with a tolerance of two positions from the annotated *ω* -site. We allow for this flexibility when calculating the F1 score as the training set contains only non-experimentally verified *ω*-site samples, which are not as reliable as the experimentally verified. The MCC is weighed twice as important as the F1 score. We weigh the MCC more as we want to emphasize the GPI-anchoring discrimination over the *ω*-site prediction performance. The model with the combination of hyperparameters that gives the best heuristics, on the validation partition, is chosen for each fold. This heuristic also controls when the model’s parameters are stored as an early stopping approach. The self evaluation during training is the Cross Entropy Loss.

#### 2.3.2. Qualitative evaluation methods

To visualize the decision making of the model, we perform a feature importance analysis using the Local Interpretable Model-agnostic Explanations (LIME) package [28]. We perform this analysis on each partition separately, using the corresponding four model ensemble. In the LIME analysis, amino acids contributing to a GPI-anchored prediction will have a positive importance, while amino acids contributing to the non GPI-anchored prediction will have a negative importance. The larger the weight, the larger the contribution to the prediction.

Furthermore, we investigate the sequence composition around the *ω*-site to uncover possible model biases.

## 3. Results and discussion

### 3.1. Quantitative results

The subset of the GPI dataset that is used for benchmarking contains 160 GPI-anchoring signal sequence samples, regarded as positive, and 2573 samples without a GPI-anchoring signal sequence, regarded as negative. To benchmark the *ω*-site position prediction the positive set is limited to the 50, out of the 160 positive protein samples, with an experimentally verified *ω*-site annotation.

To benchmark the performance of the existing tools the dataset was submitted to the three tools currently available; Big-∏, GPI-SOM, and PredGPI. In the case of Big-∏ we separated the benchmark set according to kingdom and submitted to the corresponding versions of the tool. Big-∏ annotates its predictions according to likelihood. Predictions with high likelihood are labeled as *P*, twilight zone predictions are labeled as *S*, and non-potentially GPI-anchored proteins are labeled as *N*. We regarded any protein predicted as potentially GPI-anchored (*P* or *S*) as a GPI-anchored prediction.

PredGPI ranks and classifies predictions according to specificity. Predictions are regarded as highly probable, probable, weakly probable, and not GPI-anchored. We measure the performance for two settings of PredGPI; designating weakly probable either as GPI-anchored or non GPI-anchored. Assuming weakly probable as negative predictions gives the best performance according to MCC, as shown in table 4.

**Table 4.**
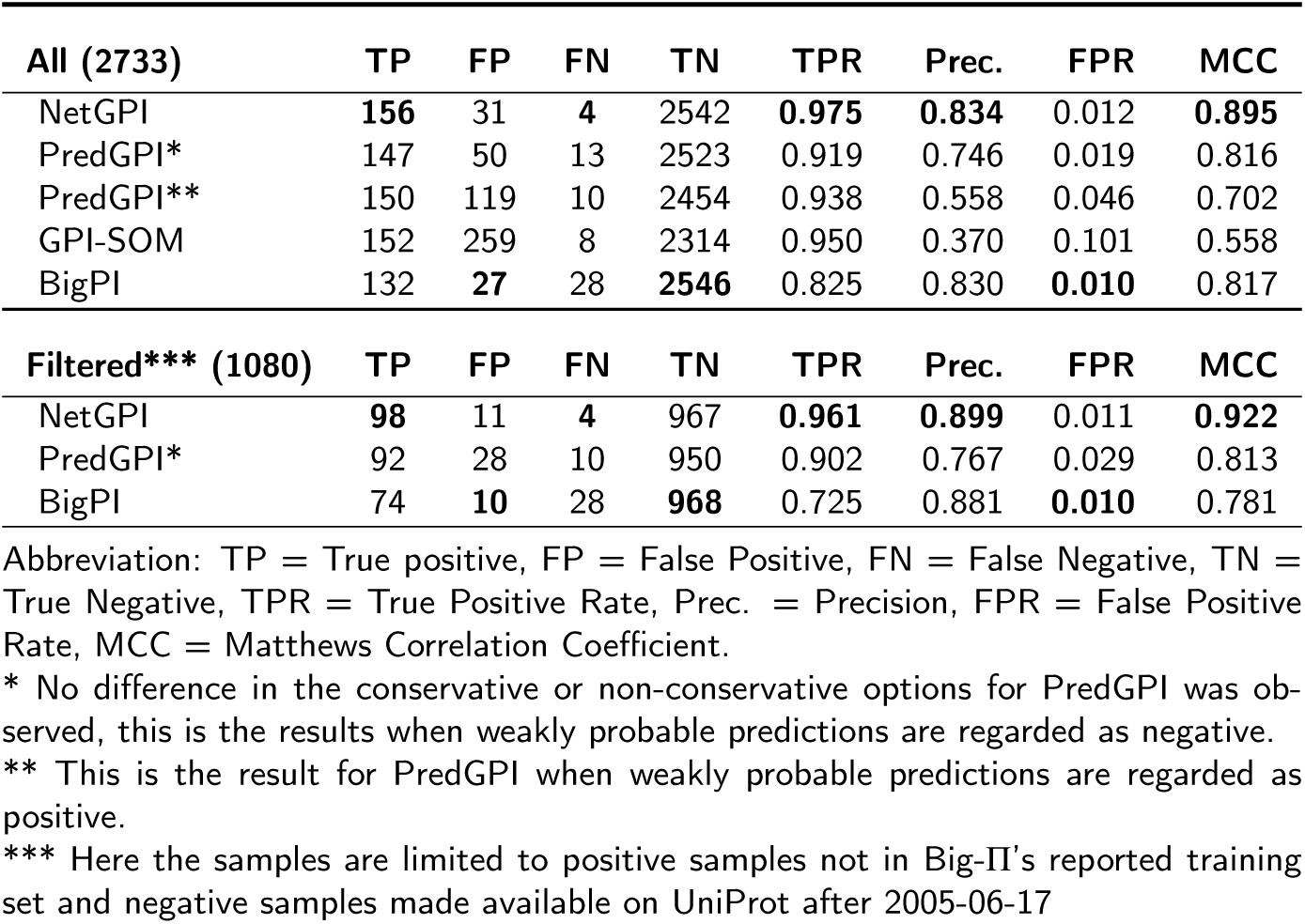
Comparison of the GPI-anchor presence prediction performance of NetGPI and benchmarked methods. NetGPI achieves superior performance on all accounts except FPR where it is outperformed by Big-∏.

For predicting the presence of GPI-anchors, NetGPI achieves the highest MCC of **0.895**. It also attains the highest true positive rate (TPR), **0.975**, the second highest being GPI-SOM. NetGPI achieves the highest precision, 0.834, the second highest being Big-∏ with a precision of 0.830. Big-∏ has the second highest MCC, 0.817 and the lowest false positive rate (FPR), 0.010, whereas NetGPI has the second lowest FPR, 0.012. For a detailed comparison see table 4.

We find that the Big-∏ learning set has at least 58 overlapping samples with our positive benchmark set and an unknown overlap with our negative benchmark set, as the negative set is not reported. This might cause the performance of Big-∏ to be overestimated. The publishing date of Eisenhaber et al. [7] is the 19th of March 2004, however the metazoa and protozoa predictors are reported to have been updated on the 17th of June 2005. We filter the benchmark set to GPI-positive samples not found in Big-∏’s reported training set and non GPI-anchored samples made available on UniProt after 2005-06-17. In the filtered comparison the performance gap between NetGPI and Big-∏ increases from 0.078 MCC to 0.141 MCC. All of Big-∏’s false negative predictions belong to the filtered dataset. If we regard PredGPI’s weakly probable as negative, the second highest MCC is, on the filtered dataset, achieved by PredGPI, with an MCC of 0.813.

For the prediction of the position of the *ω*-site we only consider the 50 proteins with an experimentally verified *ω*-site. The aforementioned dataset overlap is overly prevalent for these proteins, out of the 50 *ω*-sites, 33 are used for training the Big-∏ model.

NetGPI correctly predicts 32 out of the 50 experimentally verified *ω*-sites, with an F1 score of 0.496. NetGPI correctly predicts 8 out of the 17 not found in the reported Big-∏ training set, with an F1 score of 0.372. Big-∏ correctly predicts 38/50, with an F1 score of 0.628 and 8/17, with an F1 score of 0.457. GPI-SOM correctly predicts 9/17, however the F1 score is only 0.151 because of GPI-SOM’s higher false positive rate. If we allow for a one-off error window around the true *ω*-site, then NetGPI outperforms Big-∏, correctly predicting 44/50, with an F1 score of 0.682 and 13/17, with an F1 score of 0.605. Big-∏ correctly predicts 41/50, with an F1 score of 0.677 and 10/17, with an F1 score of 0.555. The *ω*-site position prediction results are detailed in table 5.

**Table 5.** Comparison of the *ω*-site position prediction performance of NetGPI and the benchmarked methods.

| Known*** (50) | $\pm 0$ | F1 | Sens. | Prec. | $\pm 1$ | F1 | Sens. | Prec. | $\pm 2$ | F1 | Sens. | Prec. |
| --- | --- | --- | --- | --- | --- | --- | --- | --- | --- | --- | --- | --- |
| NetGPI | 32 | 0.496 | 0.640 | 0.405 | **44** | **0.682** | **0.880** | 0.557 | **44** | **0.682** | **0.880** | 0.557 |
| PredGPI* | 29 | 0.403 | 0.580 | 0.309 | 35 | 0.486 | 0.700 | 0.372 | 36 | 0.500 | 0.720 | 0.383 |
| PredGPI | 28 | 0.389 | 0.560 | 0.298 | 36 | 0.500 | 0.720 | 0.383 | 37 | 0.514 | 0.740 | 0.394 |
| PredGPI*, ** | 29 | 0.272 | 0.580 | 0.178 | 35 | 0.329 | 0.700 | 0.215 | 36 | 0.338 | 0.720 | 0.221 |
| PredGPI** | 28 | 0.263 | 0.560 | 0.172 | 36 | 0.338 | 0.720 | 0.221 | 37 | 0.347 | 0.740 | 0.227 |
| GPI-SOM | 30 | 0.182 | 0.600 | 0.107 | 33 | 0.200 | 0.660 | 0.118 | 33 | 0.200 | 0.660 | 0.118 |
| BigPI | **38** | **0.628** | **0.760** | **0.535** | 41 | 0.677 | 0.820 | **0.577** | 41 | 0.677 | 0.820 | **0.577** |
| Known**** (17) | $\pm 0$ | F1 | Sens. | Prec. | $\pm 1$ | F1 | Sens. | Prec. | $\pm 2$ | F1 | Sens. | Prec. |
| NetGPI | 8 | 0.372 | 0.471 | 0.308 | **13** | **0.605** | **0.765** | 0.500 | **13** | **0.605** | **0.765** | 0.500 |
| PredGPI* | 5 | 0.172 | 0.294 | 0.122 | 9 | 0.311 | 0.529 | 0.220 | 10 | 0.345 | 0.588 | 0.244 |
| PredGPI | 4 | 0.138 | 0.235 | 0.098 | 9 | 0.311 | 0.529 | 0.220 | 10 | 0.345 | 0.588 | 0.244 |
| PredGPI*, ** | 5 | 0.121 | 0.294 | 0.076 | 9 | 0.216 | 0.529 | 0.136 | 10 | 0.242 | 0.588 | 0.152 |
| PredGPI** | 4 | 0.097 | 0.235 | 0.061 | 9 | 0.216 | 0.529 | 0.136 | 10 | 0.242 | 0.588 | 0.152 |
| GPI-SOM | **9** | 0.152 | **0.529** | 0.089 | 10 | 0.169 | 0.588 | 0.099 | 10 | 0.169 | 0.588 | 0.099 |
| BigPI | 8 | **0.457** | 0.421 | **0.500** | 10 | 0.555 | 0.588 | **0.526** | 10 | 0.555 | 0.588 | **0.526** |

Abbreviations: $\pm 0$ = The number of correctly predicted $\omega$ -sites, $\pm 1$ = The number of ω-site predictions within one position away from the correct position, $\pm 2$ = The number of ω-site predictions within two positions away from the correct position, F1 = f1-score, Sens. = Sensitivity, Prec. = Precision.
\* PredGPI provides two options, this is their conservative option.
\*\* This is the result for PredGPI when weakly probable predictions are regarded as positive.
\*\*\* For the position prediction we use the experimentally tested sequences with known $\omega$ -sites. The precision is calculated w.r.t. the experimentally tested sequences with known $\omega$ -sites as well as all negative samples.
\*\*\*\* Here the samples are limited to positive samples not in Big-PI's reported training set and negative samples made available on UniProt after 2005-06-17.

### 3.2. Qualitative results

In the qualitative analysis we investigate the importance of biological features when NetGPI predicts GPI-anchor presence and the *ω*-site. In addition, we analyze the *ω*-site composition to understand the neighborhood of true and predicted *ω*-site positions. Lastly, we investigate model likelihood of the predictions, and how it relates to model correctness.

#### 3.2.1. Feature Importance Analysis

Figure 2 illustrates the results of the LIME analysis for both positive (see Figure 2a) and negative (see Figure 2b) samples. We observe that the presence of a hydrophobic tail contributes the most towards a positive prediction. This is consistent with the literature [23], which defines the presence of a hydrophobic region from the position *ω* + 10. From that position the feature importance is much higher than for the rest of the sequence, which means that the main feature driving the positive prediction of NetGPI is the presence of the hydrophobic region. Regarding the negative predictions, we observe that the amino acids contributing the most towards a negative prediction are charged and polar amino acids. This indicates that the model is attributing higher importance to non-hydrophobic amino acids, indicating a lack of hydrophobic tail, when making a negative prediction.

**Figure 2:**
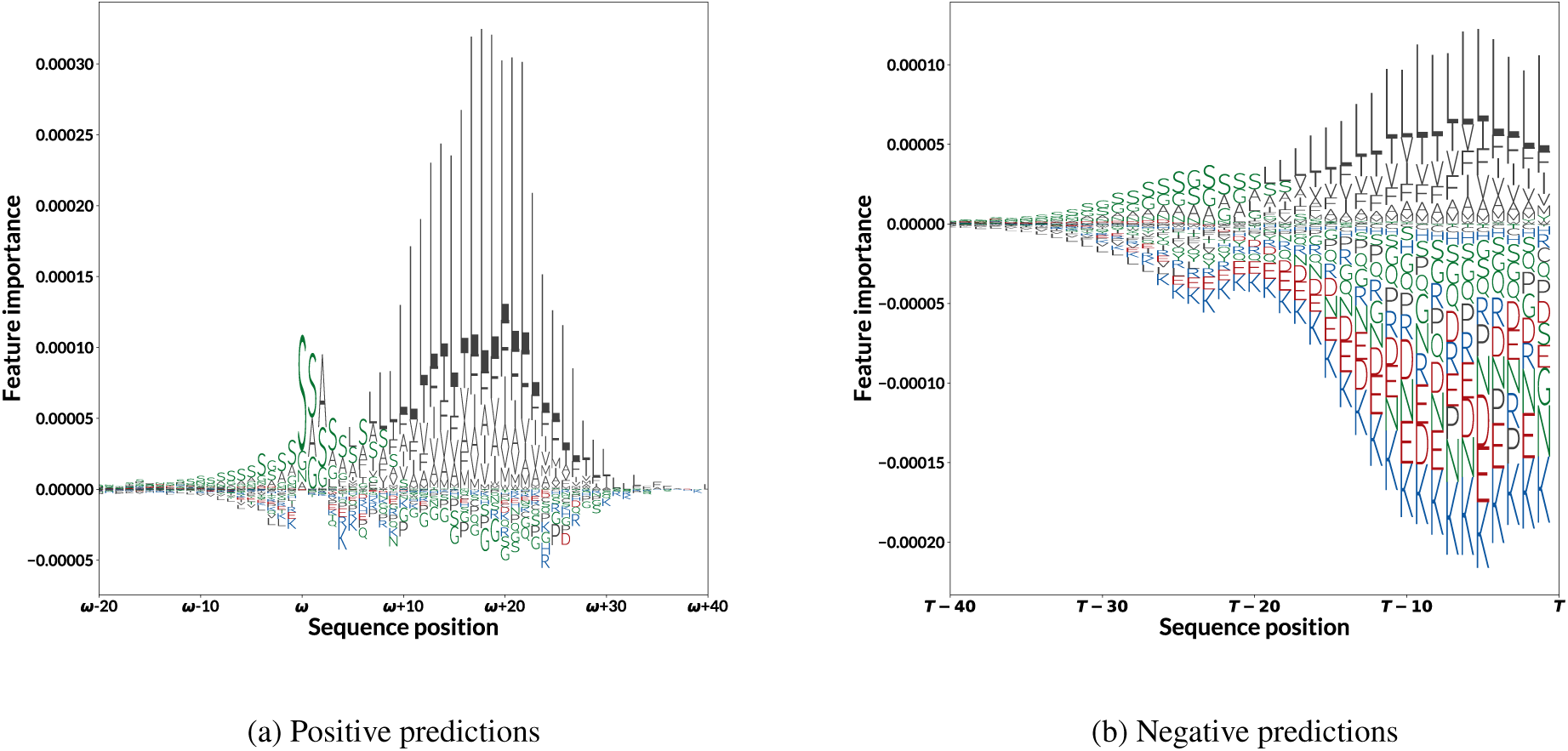
The logo plots of the LIME analysis for both positive (a) and negative (b) samples. The logo plots are colored according to amino acid properties, where blue means positively charged, green means polar, red means negatively charged and gray means hydrophobic amino acids. The positive set (a) is aligned to the predicted *ω*-site, while the negative set (b) is aligned to the C-terminus. Positive feature importance contributes to a positive prediction whereas a negative feature importance contributes to a negative one. We see that the presence of a hydrophobic tail contributes the most towards a positive prediction, whereas charged and polar amino acids contribute the most towards a negative prediction.

#### 3.2.2. ω-site composition

Out of the 50 proteins with an experimentally verified *ω*-site annotation there are 25 metazoa (animal) proteins, 13 plant proteins, 7 fungi proteins and 5 protozoa (other) proteins. Of the 25 animal proteins there are 14 which belong to *Homo sapiens*. All of the 13 plant proteins belong to the same species, *Arabidopsis thaliana*. Of the 50 experimentally verified *ω*-sites, 27 are Serine, while the other amino acids observed are Asparagine, Glycine, Aspartic acid, Cysteine and Alanine, in decreasing order of frequency. The *ω*-site of five of the *Arabidopsis thaliana* proteins are correctly positioned by NetGPI. The other eight are all one-off errors and constitute 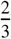 of one-off errors made by NetGPI. Out of those, there are seven where the *ω*-site amino-acid is Serine (S) where the predicted amino-acid is the Aspartic Acid (D) in the *ω*+1 position. All seven have in common the 4-mer [*ω* − 2*, ω* + 1] motif PTSD, followed either by Glycine (G) or Alanine (A) in position *ω*+2. Both Big-∏ and NetGPI are unable to position 3 out of 4 Aspartic acid *ω*-sites. This may be related to the *ω* + 2 position, as these 3 samples have a non-standard amino acid (i.e. something other than G, A, or S). See table 6.

**Table 6.**
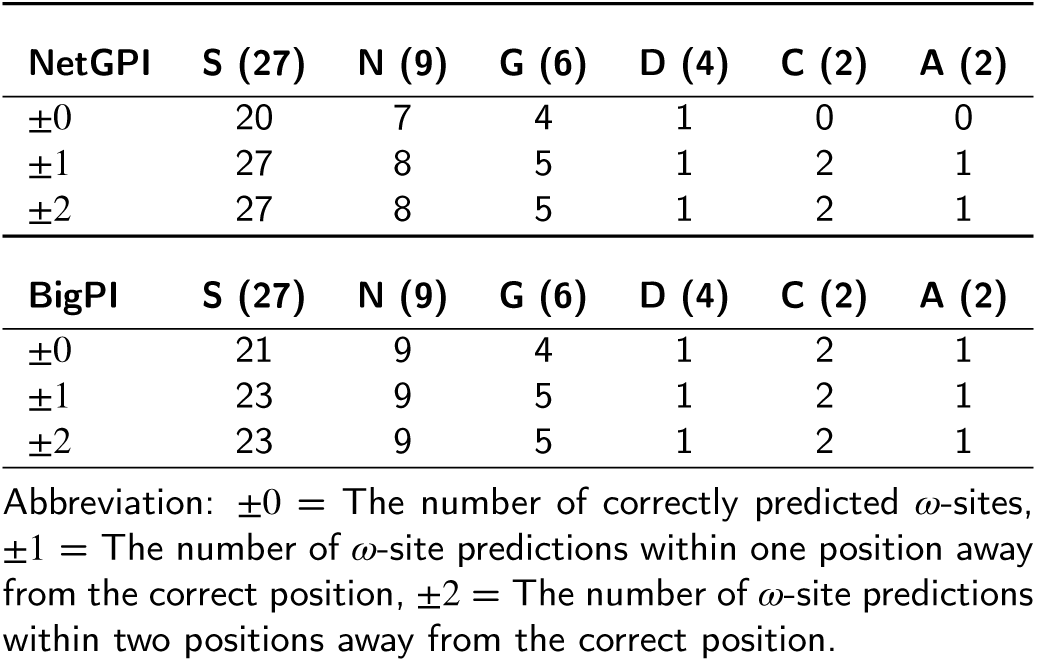
NetGPI’s and Big-∏’s *ω*-site position prediction performance for the 50 true *ω*-site amino acid in the test set. We see that both models only predict one out of four Aspartic acid *ω*-sites correctly. NetGPI has twelve one-off errors, seven of which are actually Serine *ω*-sites. The seven are homologous *Arabidopsis thaliana* proteins, which have in common the tetramer [*ω* − 2*,ω* + 1] motif PTSD, followed either by Glycine (G) or Alanine (A) in position *ω*+2.

| NetGPI | S (27) | N (9) | G (6) | D (4) | C (2) | A (2) |
| --- | --- | --- | --- | --- | --- | --- |
| $\pm 0$ | 20 | 7 | 4 | 1 | 0 | 0 |
| $\pm 1$ | 27 | 8 | 5 | 1 | 2 | 1 |
| $\pm 2$ | 27 | 8 | 5 | 1 | 2 | 1 |

| BigPI | S (27) | N (9) | G (6) | D (4) | C (2) | A (2) |
| --- | --- | --- | --- | --- | --- | --- |
| $\pm 0$ | 21 | 9 | 4 | 1 | 2 | 1 |
| $\pm 1$ | 23 | 9 | 5 | 1 | 2 | 1 |
| $\pm 2$ | 23 | 9 | 5 | 1 | 2 | 1 |

Abbreviation: $\pm 0$ = The number of correctly predicted $\omega$ -sites, $\pm 1$ = The number of ω-site predictions within one position away from the correct position, $\pm 2$ = The number of ω-site predictions within two positions away from the correct position.

#### 3.2.3. Likelihood and correctness

In addition to the classification of the sequence and the most likely position of the *ω*-site, NetGPI reports the likelihood of the chosen position. For positive predictions this is the predicted *ω*-site, while for negative predictions it is the sentinel.

As our model is trained with cross entropy, it is penalized with a logarithm of the correct prediction. If we predict incorrectly, with a very low likelihood for the correct position, the loss can be immense. We should thus expect that answers with a high likelihood are more credible.

In Figure 3 we display the likelihood distribution of the predictions. We observe differences in the likelihood of correct and incorrect predictions implying a correlation between likelihood and correctness. Furthermore, we observe higher likelihood in negative predictions than positive. This is expected as the probability distribution covers the last 100 amino acids as well as the added sentinel. Only the sentinel position denotes a negative prediction, while a positive prediction is spread across the 100 amino acid positions. This means that positive prediction likelihood has to cover all potential *ω*-site positions, while the negative prediction likelihood is limited to one position. Therefore, using the likelihood as ranking should be done separately for negative and positive results.

**Figure 3:**
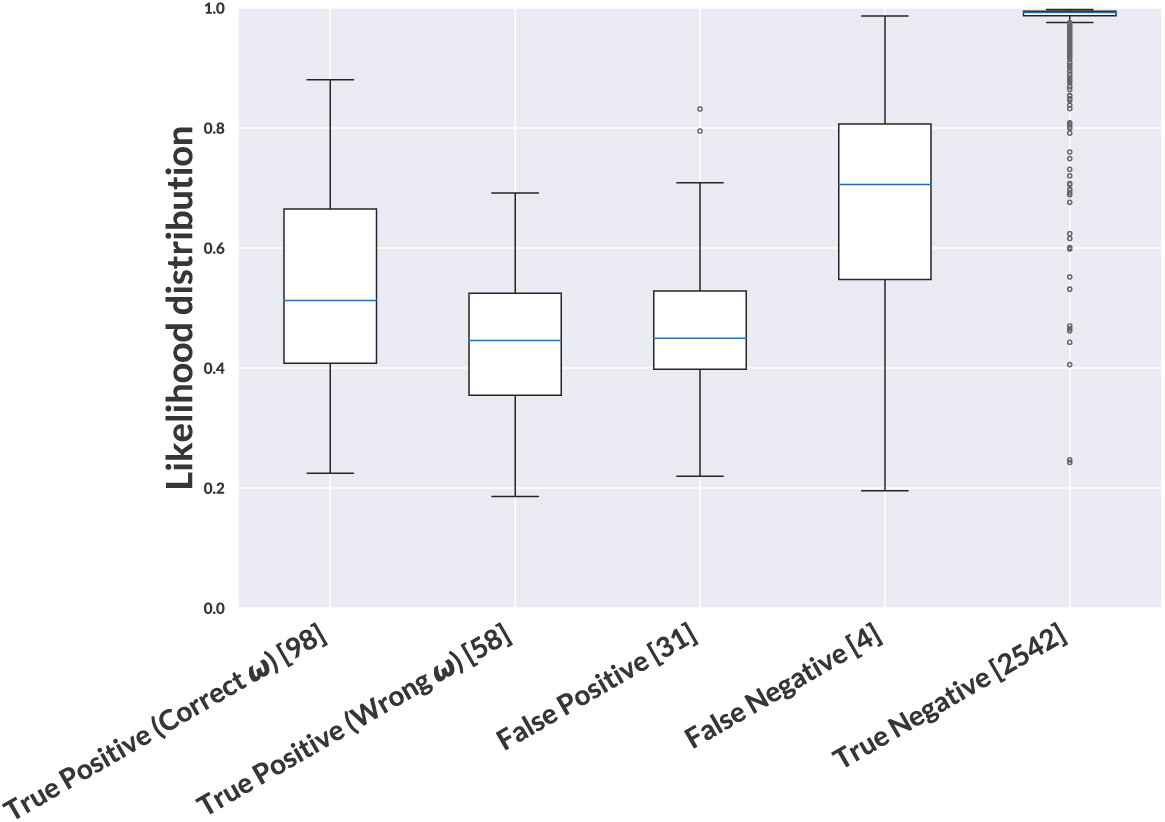
The likelihood distribution for true positive, false positive, false negative and true negative predictions. True positive are split into correctly positioned *ω*-sites and incorrectly positioned. The number of samples behind each are displayed in brackets.

## 4. Conclusion

We have shown that GPI-anchor prediction can be improved using recurrent neural networks and up-to-date datasets. Comparison with previous methods is challenging as there exists no standard dataset for training and testing predictive methods. Given progress in protein annotation, we publish a new homology partitioned dataset, using both experimentally verified proteins and manually annotated predicted proteins for training and validation. Due to the new dataset definition, the performance of current methods could be overestimated as their training sets contain sequences which are identical or homologous to sequences in our benchmark set.

Our results indicate that proteins manually annotated by prediction methods or sequence similarity are useful for training a GPI-anchor predictor to perform well when evaluated on experimentally verified GPI-anchoring signals. However, using these data may have increased the number of *ω*-site predictions that are off by one position. We believe that this limitation is necessary in order to obtain a larger training set. If we were to use only the experimentally verified GPI-anchors to train and test the predictor, we would not have enough training samples to teach a deep neural network classifier.

A web server implementing NetGPI is available at https://services.healthtech.dtu.dk/service.php?NetGPI, our dataset can be downloaded from the same site.

## Notes

https://services.healthtech.dtu.dk/service.php?NetGPI

### Competing Interest Statement

The authors have declared no competing interest.

### Summary of Updates

The version of the software has been updated to 1.1. The method for homology partitioning of the data set has been modified to ensure independence between partitions, and the neural networks have been retrained.

